# Biotic and abiotic stress distinctly drive the phyllosphere microbial community structure

**DOI:** 10.1101/2022.10.14.512112

**Authors:** Rishi Bhandari, Alvaro Sanz Saez, Courtney P. Leisner, Neha Potnis

## Abstract

While the physiological and transcriptional response of the host to biotic and abiotic stresses have been intensely studied, little is known about the resilience of associated microbiomes and their contribution towards tolerance to these stresses. We evaluated the impact of one such abiotic stress, elevated tropospheric ozone (O_3_), under open-top chamber field conditions on host susceptibility and phyllosphere microbiome associated with pepper cultivars resistant and susceptible to *Xanthomonas*. Pathogen challenge resulted in distinct microbial community structures in both cultivars under an ambient environment. Elevated O_3_ alone affected microbial community structure associated with resistant cultivar but not the susceptible cultivar, indicating the role of host genotypic background in response to abiotic stress. Elevated O_3_ did not influence overall host susceptibility but did increase disease severity on the resistant cultivar, indicating a possible compromise in the resistance. Interestingly, combined stress resulted in a shift in microbial composition and structure like that observed with pathogen challenge alone. It indicates the possible prioritization of community response towards the most significant stress and pathogen being most influential regardless of the cultivar. Despite community composition differences, overall functional redundancy was observed in the phyllosphere community. To gain insights into community-level interactions, network topology assessment indicated a stable network with enhanced taxon connectedness upon pathogen challenge. However, an observation of destabilized random network with a shift in hub taxa in the presence of combined stress warrants future studies on the consequences of such unstable microbial communities on host response to pathogens in the face of climate change.

## Introduction

The phyllosphere (aboveground parts) of plants is a unique, nutrient-poor habitat and is inhabited by various prokaryotic and eukaryotic microorganisms (1) that colonize either the leaf surface (epiphytes) or inside the leaf tissue (endophytes) (2,3). Despite being exposed to environmental fluctuations that can add stochasticity, microbial community assembly and succession processes in the phyllosphere are not considered random. Although dispersal from neighboring plants and other demographic factors like neighbor identity and age are contributing factors toward phyllosphere microbiome diversity (4), plant host factors such as host genotype, developmental stage (5), and host resistance (6,7) impose control over microbiome assembly. This host filtering of the microbiome is observed due to different resource availability on the leaf surface (8), differing physical properties (9), and host defense signaling (6,10).

Members of the phyllosphere microbiome play a vital role in maintaining plant growth and productivity (11,12). Recent studies have shown that plants can employ the “cry for help” strategy to enhance their ability to combat biotic and abiotic stresses by recruiting beneficial microbes or traits (13). Pathogen invasion is one of the most influential biotic stresses affecting the plant microbial assembly in the phyllosphere (14), where pathogens can cause a damaging alteration of the habitat by various virulence factors or through niche or resource competition (1). The increased susceptibility of the host leads to colonization by other opportunistic microbes that might not be able to cause disease alone (15). This indicates that pathogen invasion can cause a shift in the phyllosphere microbiome assembly and structure (14). Plant defense signaling activated in response to pathogen attack is also a source of disturbance in the phyllosphere community. Yet, dominant community members are thought to restore stability to this disturbed community (16). Increasing evidence has suggested that plants can recruit disease-suppressive microbes in response to pathogen attacks in the phyllosphere (17,18), similar to what has been observed in the rhizosphere (19).

Similar to biotic stress, different phyllosphere microbes are known to provide resistance against abiotic stresses such as drought (20), ultraviolet radiation (21), and various air pollutants (22). With climate change, it is predicted that plant diseases can extend beyond their geographical range, and their interactions with different or alternate hosts can change the prevalence of the diseases depending on the direction of the interacting effects (23). Besides changes in pathogen biology and epidemiology, abiotic stresses resulting from changing climate can alter host susceptibility by interfering with defense hormone networks (24) and thus influence disease incidence. Exposure of plants to both abiotic and biotic stress conditions can result in positive or negative impacts on plant responses depending on the timing, nature, and severity of each stress, as different defense signaling pathways may interact or inhibit each other (25). Global warming caused by greenhouse gases has resulted in the increase of tropospheric ozone (O_3_) due to the rise in precursors such as nitrogen oxide (NOx), CO, methane, and other volatile organic compounds (26,27). A study across the US predicted that the 95^th^ percentile for daily 8-hour maximum O_3_ will increase from 31-79 parts per billion (ppb) in 2012 to 30-87 ppb in 2050 (28). As O_3_ is highly phytotoxic, it is reported to affect plants in a variety of ways, including visible injury and reduction in photosynthesis which in turn affects plant growth, nutritional value, crop yield, and alterations to carbon allocation, and has the potential to affect the plant microbiome (29–31). A contrasting influence of elevated O_3_ on plants in the presence of various pathogens has been observed, indicating that the effect of O_3_ on disease severity does not depend on the pathogen’s lifestyle but the interaction between the host, environment, and pathogen (32–35).

While several studies have focused on understanding how plants respond to a combination of biotic and abiotic stress at the physiological, transcriptional, and cellular levels (36), how plants manipulate their microbiome while responding to a combination of stresses has not been explored. In this study, our overall goal is to obtain a comprehensive view of plant resilience to the future climate in the presence of biotic stress. We explicitly focused on the response of the phyllosphere microbiome to individual and combined biotic and abiotic stresses and dissected the influence of genotype x environment (G x E) interactions on the overall outcome of plant disease as well as on microbiome structure and function. Our study addressed these knowledge gaps using a field experiment involving open-top chambers (OTCs) that allowed us to manipulate O_3_ levels. We investigated the structure, seasonal succession, function, and network of phyllosphere microbial communities associated with two pepper cultivars (resistant and susceptible to infection by a pathogen, *Xanthomonas perforans*) grown under two environmental conditions (ambient and elevated O_3_) in the presence and absence of a pathogen (inoculated and control). We hypothesized that host resistance would be compromised due to an increase in O_3_ stress (here referred to as abiotic stress), and there would be functional redundancy and resilience among the phyllosphere microbiome in response to biotic (infection with *Xanthomonas*) and abiotic stress. We also hypothesized that under the influence of combined stress (elevated O_3_ and infection with *Xanthomonas*), the plant would prioritize its response towards the most impactful stress, eventually driving the microbial community structure.

## Materials and Methods

### Experimental site and design

The experiment was conducted at the Atmospheric Deposition (AtDep) site at Auburn University (Fig. S1A) in the 2021 growing season (May-July), where we harnessed OTCs (Fig. S1B) that allowed us to test the effect of O_3_ stress on plant-pathogen-microbiome interactions and address the complexity of plant defense-development trade-off. We used 12 chambers for fumigation, where six chambers had an ambient environment, and six had elevated O_3_. Each elevated O_3_ chamber contains four O_3_ generators (HVAC-1100 Ozone generator, Ozone Technologies, Hull, IA, USA), and each generator powers two ultraviolet (UV) bulbs (Model GPH380T5VH/HO/4P, Ozone Technologies, Hull, IA, USA). Generators and bulbs are located within the elevated O_3_ chamber fan boxes. To reach the desired set-point (∼100 ppb), O_3_ generators were controlled by 0-10V control wires, which are controlled via an analog output module. Each chamber is connected via plastic tubes to a central gas manifold to which each chamber is opened sequentially by 3-way solenoid valves. To fumigate the plants, the ozonated air was blown from the fan box into the plastic lining of the open-top chamber (Fig. S1B). The plastic panel on the portion of the chamber is double-walled with holes on the inside panel, allowing O_3_ to be released over the plants inside the chamber. A microcontroller cycles through the 12 solenoid valves every 24 minutes (sampling each of the 12 chambers for 2 minutes) to monitor O_3_ from each chamber (Model 205 Dual Beam Ozone Monitor, 2B Technologies, Boulder, CO, USA) during the fumigation window (10 am to 6 pm) to sample the O_3_ concentration in each chamber (both ambient and elevated). In each of the chambers, we had six plants, each of Early California Wonder (referred to hereafter as susceptible cultivar) and PS 09979325 (referred hereafter as resistant cultivar). Among the 12 chambers, plants in 6 chambers were inoculated with the pathogen *Xanthomonas perforans*. During this experiment, the average [O_3_] in the control chambers was around 30.6 ppb, while the fumigated chambers had an average [O_3_] of about 90.3 ppb (Fig. S1C).

### Disease severity and pathogen population

The effect of host resistance and presence of abiotic stress on overall disease development and pathogen population was evaluated by dip inoculating 5–6-weeks old pepper plants from both cultivars with *Xanthomonas perforans* suspension adjusted to 10^6^ CFU/ml in MgSO_4_ buffer amended with 0.0045% (vol/vol) Silwet L-77 (PhytoTechnology Laboratories, Shawnee Mission, KS, USA). The control plants were dip-inoculated in MgSO_4_ buffer. The dip-inoculated plants were transferred to the above-mentioned OTCs. Disease severity assessments were made by estimating the percentage of disease symptoms and defoliation caused by bacterial spots using the Horsfall-Barratt scale (37) during both the mid and end season, and the pathogen abundance was correlated based on the relative abundance of *Xanthomonas* in each treatment during both mid and end season.

### Sample collection, DNA extraction, sequencing, and quality trimming

Pepper leaf samples were collected from inoculated and control samples of each cultivar before keeping the plants in the chambers (base samples), followed by two other time points during the growing season (mid and end season). The leaf samples were then used to extract and quantify the metagenomic DNA as described earlier (38). The DNA samples were submitted to the Duke Center for Genomic and Computational Biology sequencing core (Duke University, Durham, NC) for library preparation, and paired-end reads (2 × 150 bp) were sequenced on NovaSeq 6000 S1 flow cell system. The raw reads were then trimmed for quality and host contamination, as described earlier (38).

### Taxonomic profiling

Quality controlled and host decontaminated reads were taxonomically assigned using Kraken2 (v2.1.2) (39) against a standard Kraken2 database containing RefSeq libraries of (40) of archaeal, bacterial, human, and viral sequences (as of 03/01/2022). Kraken2 is a kmer based short read classification system that assigns a taxonomic identification to each sequencing read by using the lowest common ancestor (LCA) of matching genomes in the database. Kraken2 report files were used as inputs to run Bayesian re-estimation of abundance with the Kraken (Bracken) (v2.6.2) (41) to re-estimate abundance at each taxonomic rank for all the samples. Bracken uses the taxonomy labels assigned by Kraken to estimate the abundance of each species. The database for Bracken was subsequently built with the Kraken2 database using the default 35 k-mer length and 100 bp read lengths based on the average read length in our sample with the lowest read length to re-estimate the abundance of microbial communities at the species level. The outputs from Bracken were combined using the combine_bracken_outputs.py function for downstream analysis. The kraken-biom tool (https://github.com/smdabdoub/kraken-biom) was used to convert the output from Bracken into BIOM format tables for diversity analyses in R (42).

The taxonomic composition and diversity of eukaryotes in the samples were accessed using the EukDetect (43). EukDetect aligns the metagenomic reads to universal marker genes from conserved gene families curated from fungi, protists, non-vertebrate metazoan, and non-streptophyte archaeplastida genomes and transcriptomes followed by low-quality and duplicate reads filtering. The final eukaryote abundance is calculated by filtering taxa with fewer than four reads and aligning to less than two marker genes. The resulting absolute abundance (Reads Per Kilobase of Sequence) was used to compare the diversity across the samples. The RPKS value was normalized by multiplying with a scaling factor calculated by dividing the median library size by the sample library size, which was then used to compare across the samples.

### Diversity, statistical, and network analysis

All statistical and diversity analyses were performed using R (v 4.1.3) (42) and Rstudio (44) with the packages Phyloseq (v1.38.0) (45), *vegan* (v2.5-7) (46), *ggplot2* (v 3.3.5) (47) packages. Before data analysis, the library size was normalized using scaling with ranked subsampling with ‘SRS’-function in the ‘SRS’ R-package (v 0.2.2) (48). Alpha diversity measures Chao1 and Shannon index were used to identify community richness and diversity, respectively. The Wilcoxon rank sum test tested significant differences in alpha diversity indices for nonparametric data and the T-test for normally distributed data. The appropriateness of these methods was verified by checking for the normal distribution of residuals based on the Shapiro-Wilk test for normality.

Furthermore, the differences in overall microbial profiles among the cultivars and different environmental conditions (β-diversity) were estimated using the Bray–Curtis distance. To understand the factors contributing to microbial community structure, we performed permutation multivariate analysis of variance (PERMANOVA) (49) and analysis of similarities (ANOSIM) with 1000 permutations (*p* = 0.05) with different factors (cultivars, environment, inoculation status, and time of sampling) and Bray-Curtis dissimilarity. In addition, multivariate homogeneity of group dispersion test (BETADISPER) (50) was performed to determine the homogenous dispersion between the factors in relation to their microbial taxa. Non-metric multidimensional scaling (NMDS) among the sample groups was calculated using Bray-Curtis dissimilarity and visualized using the ggplot2 package in R.

For our network analysis, the taxonomic data was subsetted to at least 0.5% relative abundance in over 20% of the samples (prevalence). Correlation network analysis was performed using the SPRING (51) approach implemented in the R package NetCoMi (52). Community structures across the treatment were estimated using the “cluster_fast_greedy” algorithm (53), and hub taxa were determined using the threshold of 0.95. A quantitative network assessment was performed with a permutation approach (1000 bootstraps) with an adaptive Benjamini-Hochberg correction for multiple testing.

### Functional profiling and comparison

Functional profiling of the microbial communities was conducted on concatenated paired-end sequences with HUMAnN3 (v3.0) (54). ChocoPhlAn nucleotide database v30 was used for functional pathway abundance and coverage estimation. The resulting pathway abundance tables were sum-normalized to copies per million reads (CPM) to facilitate comparisons between samples with different sequencing depths. The output from HUMAnN3 was then imported into QIIME2 (55) to calculate different diversity indices. These calculated diversity indices were then analyzed using the vegan (v2.5-7) package in R (v 4.1.3) to test the overall diversity in functional profiles. Differentially abundant pathways across the treatment were identified using the LEfSe (version 1.1.2) (56). Pathways with a corrected *p*-value of 0.05 or less were classified as significantly increased within one of the two groups.

## Results

### Elevated O_3_ did not influence disease severity levels on a susceptible cultivar but did support higher disease severity on a resistant cultivar compared to the ambient environment

Disease severity measured by the percentage of bacterial spot disease symptoms and defoliation was recorded on the susceptible and resistant cultivar during the mid and end season, both under ambient and elevated O_3_. As expected, higher disease severity values were recorded on susceptible cultivars compared to resistant cultivars. Elevated O_3_ did not impact disease severity on susceptible cultivar. However, significantly higher disease severity levels were observed on resistant cultivars under elevated O_3_ conditions, both at mid-season (*p* < 0.001) and end season (*p* = 0.01) (Fig. 1, Table S1), indicating possible compromise in disease resistance against *Xanthomonas* under O_3_ stress.

**Fig. 1.**
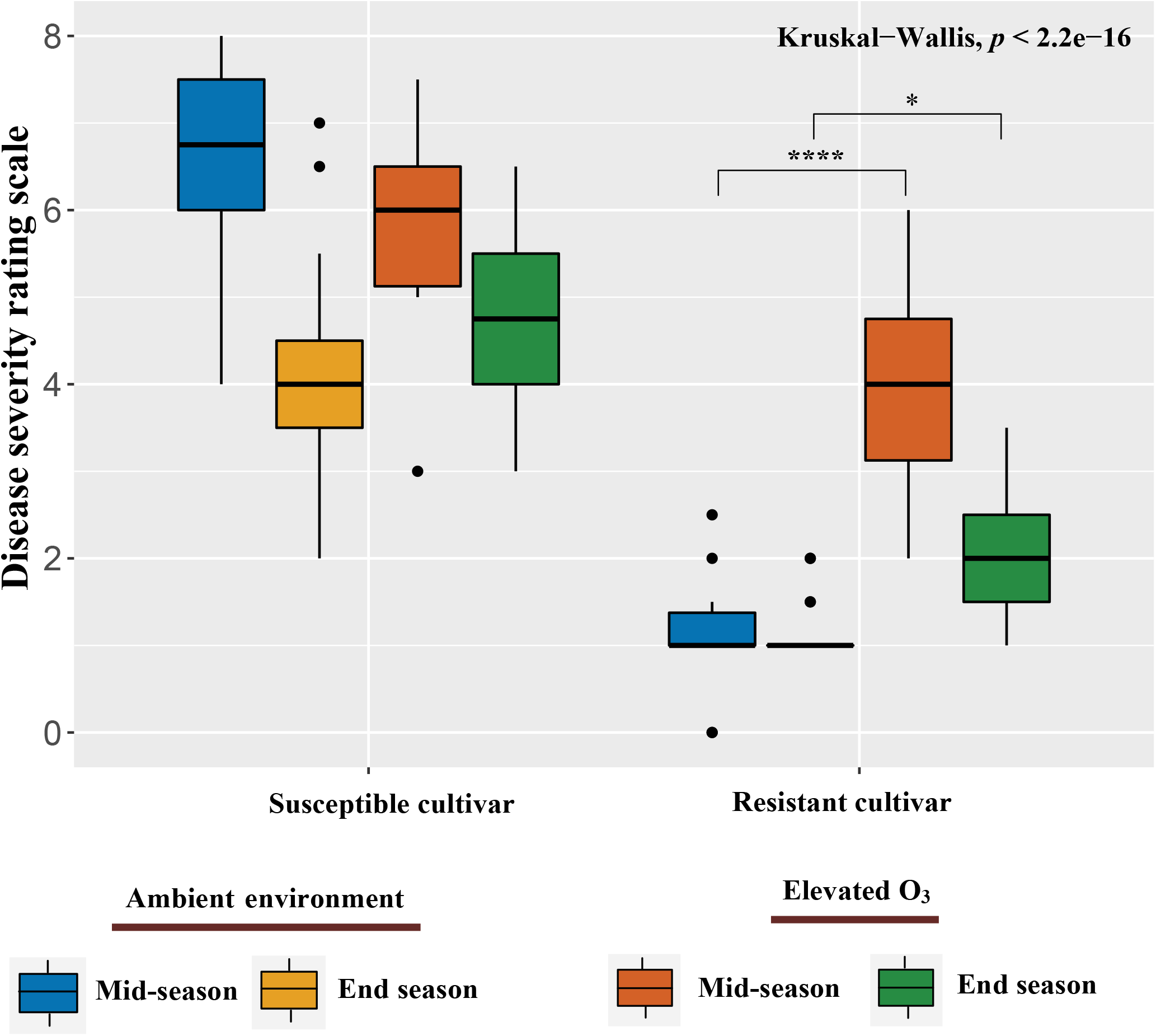
Box and whisker plots showing the disease severity under elevated O_3_ and ambient environmental conditions across susceptible and resistant cultivars. Significance levels for each of the treatment combination are indicated by \**p* < 0.05; \*\**p* < 0.01; \*\*\**p* < 0.001; \*\*\*\**p* < 0.0001.

### Sequencing statistics

The samples collected twice during the growing season were subjected to shotgun metagenome sequencing, which produced 2.83 to 17.16 Gbps of raw reads per sample. Adapter trimming and removal of low-quality reads resulted in the loss of 4.3 to 11.3% of the total reads among different samples. Of the quality trimmed reads, 5.78 to 39.09% of the reads were identified as host reads and removed from further analysis. The samples at the early seedling stage yielded very few reads upon filtering because of higher host contamination (23-39%), indicating minimal microbial colonization in the greenhouse-grown seedlings before transplanting. Around 50.61% to 84.56% of the original total reads were retained for downstream analysis (Table S2).

### Host susceptibility affects microbial diversity and richness

We next investigated the effect of biotic and abiotic stress and their interaction on overall microbial diversity and richness of phyllosphere communities. Overall richness and diversity values in both the mid and end season samples were higher than baseline samples. This could be attributed to low microbial colonization levels on greenhouse-grown base samples that increased in diversity and richness upon exposure to natural field conditions. Inoculation, time of sampling during the growing season, or elevated O_3_ did not influence bacterial richness (*p* = 0.086) (Fig. 2A) and diversity (Kruskal-Wallis, *p* = 0.055) (Fig. 2B), or eukaryotic richness (Kruskal-Wallis, *p* = 0.087) (Fig. 2C) and diversity (Kruskal-Wallis, *p* = 0.23) (Fig. 2D) for the resistant cultivar. However, inoculation with *Xanthomonas* led to significantly lower bacterial richness (*p* < 0.001) (Fig. 2A) and diversity (Kruskal-Wallis, *p* = 0.014) (Fig. 2B) as well as lower eukaryotic richness (Kruskal-Wallis, *p* = 0.013) (Fig. 2C) and diversity (Kruskal-Wallis, *p* = 0.021) (Fig. 2D) for the susceptible cultivar under ambient conditions throughout the growing season. The O_3_ levels alone did not influence bacterial (Table S3) and eukaryotic richness and diversity (Table S4), either in the presence or absence of a pathogen in both cultivars.

**Fig. 2.**
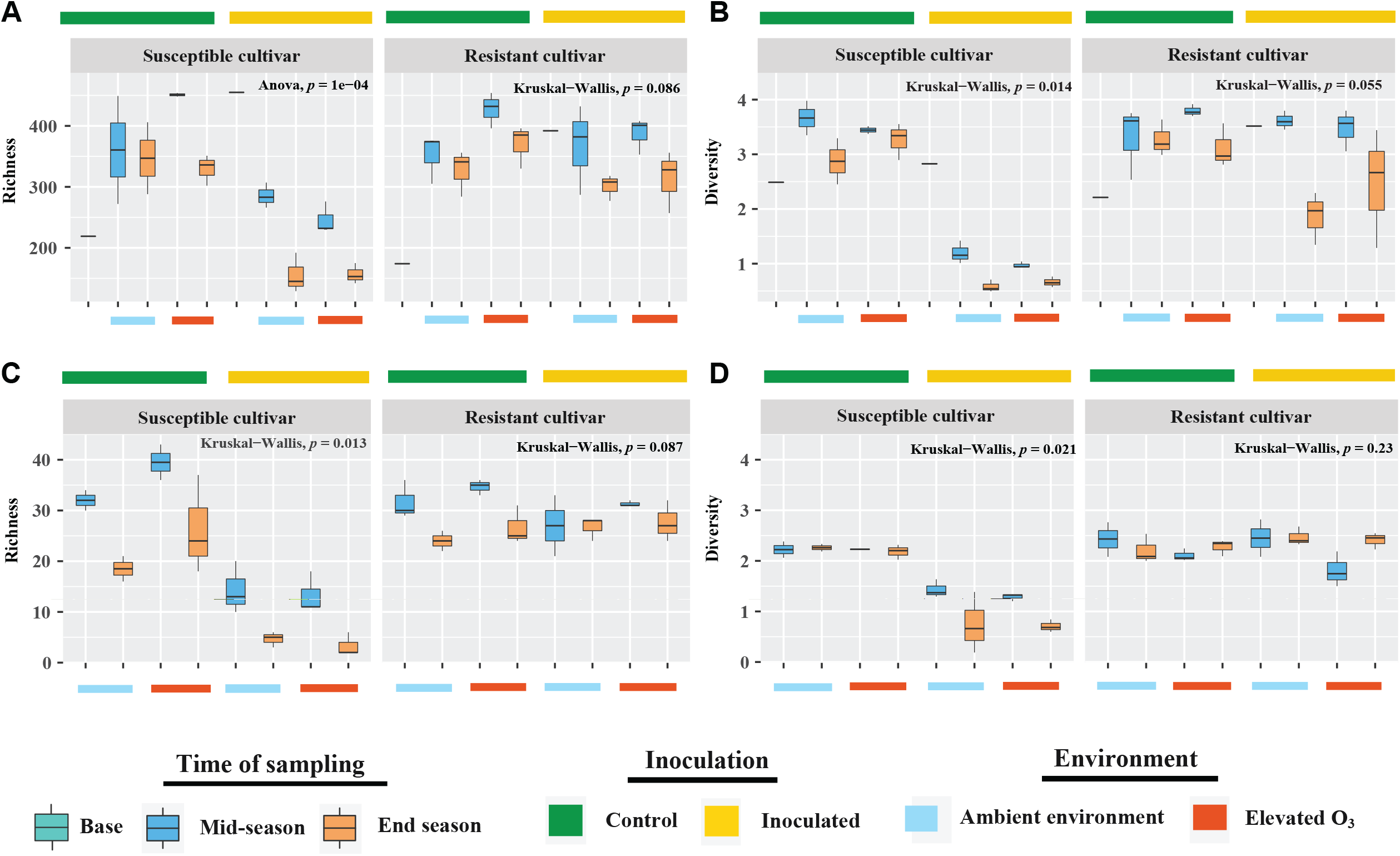
Box and whisker plot showing microbial community diversity and richness across different treatment conditions in susceptible and resistant cultivars. (A) Bacterial Chao1 richness and (B) bacterial Shannon diversity index across different environments. (C) Eukaryotic community diversity and (D) richness across different treatments. Inoculated and control samples are indicated with yellow and green bars on the top, while ambient and elevated ozone treatments are denoted by light blue and red color bars at the bottom, respectively.

### The effect of O_3_ levels was significant on the eukaryote community, yet was minimal in shaping bacterial community structure

To visualize the differences in bacterial and eukaryotic community structure between samples from two pepper genotypes and two environmental conditions, the taxonomic abundance profiles were used to compute the Bray-Curtis distance matrix and plotted into two dimensions using non-metric multidimensional scaling (NMDS). Bacterial communities associated with inoculated susceptible cultivar clustered distinctly from those associated with control susceptible as well as inoculated and control resistant cultivar. We further observed distinct clusters separating bacterial communities based on temporal sampling (Fig. 3A). Regardless of the inoculation status, there was a separation between eukaryotes communities recovered from susceptible and resistant cultivars. The influence of temporal sampling on clustering was evident in eukaryotes communities like those observed in bacterial communities (Fig. 3B).

**Fig. 3.**
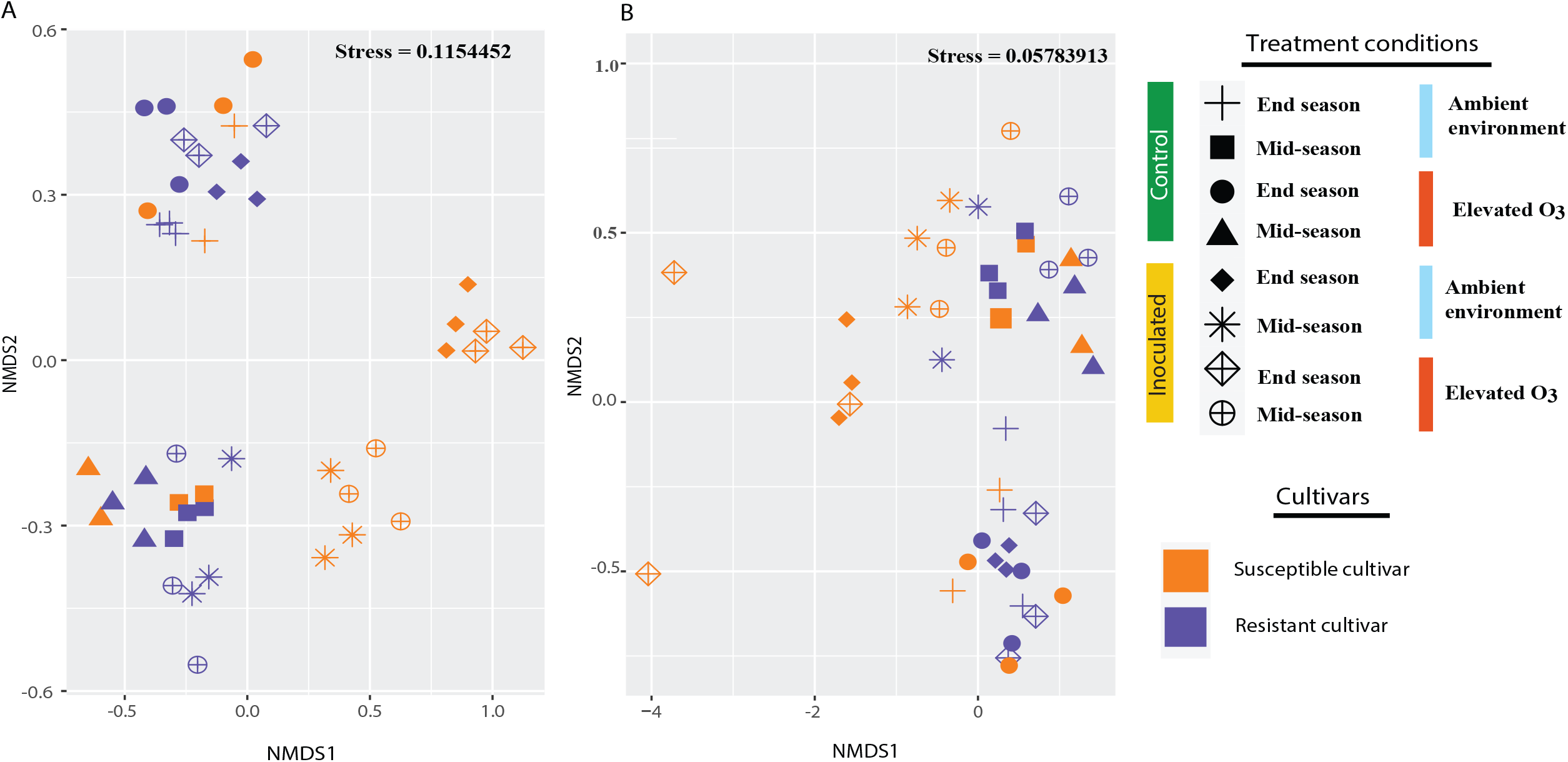
Nonmetric Multidimensional Scaling (NMDS) ordination displaying diversity in community composition across different treatment conditions. (A) NMDS ordination comparing the bacterial community diversity across two cultivars, environmental conditions, and time of sampling. (B) NMDS ordination comparing the eukaryotic community diversity across two cultivars, environmental conditions, and time of sampling.

To understand the relative influence of each factor and their interaction on the overall phyllosphere microbial community structure, we performed a PERMANOVA on Bray-Curtis dissimilarities using cultivar, time of sampling, environment, and inoculation as independent variables. Overall, the effect of cultivar, time of sampling, and inoculation were highly significant in shaping bacterial communities (*p* < 0.001) in addition to the interactions of cultivar, time, and inoculation *(p* = 0.032) (Table S5A). We further assessed individual factors’ influence and interactions across two sampling points. The effect of the cultivar was significant but diminished over the growing season (mid-season: R^2^ = 0.232, *p* < 0.001; end season: R^2^ = 0.064, *p* = 0.032). In contrast, effect of inoculation increased over the course of growing season (mid-season: R^2^ = 0.209, *p* < 0.001; end season: R^2^ = 0.553, *p* < 0.001) (Table S5B, Table S5C). The effect of interaction among cultivar and inoculation on bacterial communities remained statistically significant over time, although the effect decreased in size by the end of the growing season (mid-season: R^2^ = 0.157, *p* = 0.001; end season: R^2^ = 0.052, *p* = 0.042). The effect of the environment was minimal, with it being not statistically significant by the end of the growing season (mid-season: R^2^ = 0.054, *p* = 0.048; end season: R^2^ = 0.028, *p* = 0.152) (Table S5B, Table S5C). The interaction between the environment and other variables was insignificant throughout the growing season. An increase in O_3_ levels did not alter the bacterial community structure on the susceptible cultivar. However, it did influence bacterial communities on the resistant cultivar (R^2^ = 0.141, *p* = 0.027) (Table S5D) in the absence of *Xanthomonas*.

Like bacterial communities, eukaryote diversity was also significantly influenced by the environment, cultivar, time of sampling, and inoculation (*p* < 0.01) (Table S6A, Table S6B). Cultivar had a significant effect on eukaryote diversity with more influence during the end season (mid-season: R^2^ = 0.120 (Table S6C), *p* = 0.007; end season: R^2^ = 0.374, *p* = 0.001 (Table S6D, Table S6E)). An increase in O_3_ level significantly affected the eukaryotic community during the mid-season, while it was not significant during the end season (mid-season: R^2^ = 0.222, *p* = 0.001 (Table S6C); end season: R^2^ = 0.066, *p* = 0.19 (Table S6D, Table S6E). The effect of inoculation on eukaryotic communities was higher during the mid-season, and it decreased during the end season (mid-season: R^2^ = 0.154, *p* = 0.003 (Table S6C); end season: R^2^ = 0.115, *p* = 0.035 (Table S6D, Table S6E)).

These findings indicate that distinct phyllosphere microbial communities were found between the resistant and susceptible cultivar, which in turn, is differentially affected by sampling time, the presence of a dominant pathogen, abiotic stress, and the interaction between these factors. Among these factors, cultivar had more influence in shaping the bacterial communities during the early time, while inoculation had more impact later in the season. Regarding eukaryotes, diversity was primarily influenced by elevated O_3_ during the mid-season, while host genotype had more effect during the end season.

### Phyllosphere microbial taxa are differentially affected by biotic and abiotic stress during the growing season

We characterized the microbial community on resistant and susceptible cultivars in the presence of elevated O_3_ and ambient environmental conditions to investigate the temporal dynamics in community assembly and succession in the phyllosphere, and patterns were compared between inoculated and control plants. Detailed taxonomic description of bacteria and eukaryotes across different treatments is given in supplementary information. The taxonomic diversity analysis showed that several bacterial (Fig. 4A, Table S7) and eukaryotic genera (Fig. 4B, Table S8) monopolizing the phyllosphere environment. These microbial genera are differentially affected by the presence of a pathogen, environmental stress, and their interaction.

**Fig. 4.**
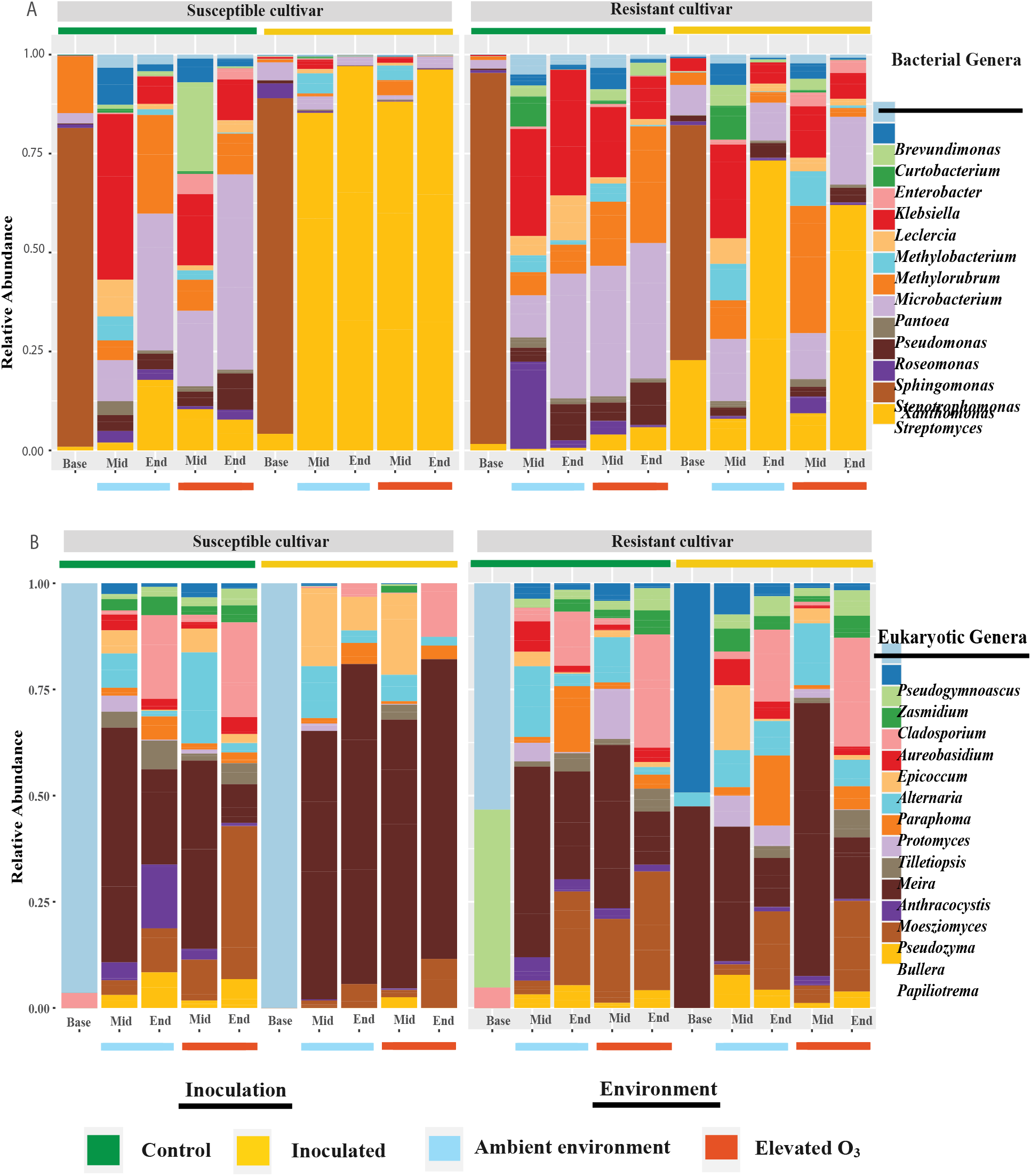
Stacked bar plot showing the relative abundance of genus-level taxonomic profiles across the samples for susceptible and resistant cultivars. (A) Bar plots showing the relative abundances of the top 15 bacterial genera across the samples. (B) Bar plots showing the relative abundance of the top 15 eukaryotic genera across the samples. Control samples are indicated by a green bar, while the yellow bar represents the inoculated samples. The time of sampling is indicated by Base (initial samples), Mid (mid-season), and End (end of the season). Ambient represents the normal environment, whereas elevated O_3_ represents the chambers with elevated O_3_ in the treatment.

Taxonomic diversity analysis of *Xanthomonas* indicates the presence of *Xanthomonas* on control plants of susceptible and resistant cultivars, suggesting low levels of natural inoculum in the field. However, the relative abundance of *Xanthomonas* on control plants did not increase significantly over time (< 5% by the end season). As expected, the relative abundance of *Xanthomonas* increased from 86 % during the mid-season to 96 % end season on inoculated susceptible cultivar and 8 % during the mid-season to 66% during the end season on resistant cultivars. Significant variation in the relative abundance values for the *Xanthomonas* population during the end season (∼33-96%) on resistant inoculated leaves under elevated O_3_ conditions was worth noting. However, elevated O_3_ did not reveal a significant difference in the *Xanthomonas* population in either cultivar (Fig. S4). This observation was surprising given that disease severity levels under elevated O_3_ conditions on resistant inoculated plants were significantly higher than that under ambient conditions, suggesting that the abundance of the pathogen population is not the determinant of the disease severity levels under elevated O_3_.

In susceptible cultivar, bacterial genera *Pseudomonas, Pantoea, Methylobacterium, Sphingomonas, Methylobrum*, etc., were negatively affected, while *Microbacterium* was positively influenced in the presence of *Xanthomonas*. In contrast to the susceptible cultivar, the relative abundance of *Pseudomonas* and *Sphingomonas* increased in the presence of the *Xanthomonas* in the resistance cultivar. The bacterial genus *Methylobacterium* was negatively influenced by elevated O_3_, while the genera *Pseudomonas* and *Sphingomonas* were positively impacted. Regarding eukaryotes, the genus *Bullera* was positively affected by elevated O_3_, while the genus *Epicoccum* and *Protomyces* had temporal variation regardless of treatment.

### Phyllosphere microbial communities with different taxonomical compositions show functional redundancy

Although the microbial composition was significantly affected by cultivar, inoculation, and time, overall metabolic pathways based on the abundance of microbial metabolic pathways and associated functions in each community remained insignificant (*p* > 0.05) (Fig. S5, Table S9A). The result suggested that communities encode functional redundancy despite their distinct taxonomic structure. However, the interaction between inoculation, cultivar, and sampling time had a significant effect on microbial functions (*p* = 0.001). Assessing the functional diversity across the time points, we observed the considerable effect of the cultivar during the end season (*p* = 0.019) (Table S9B). In contrast, all other factors remain insignificant during both time points. We observed substantial variability at the functional level across all groups when looking closely at individual metabolic pathways enriched across treatment comparison. Inoculation with *Xanthomonas* altered microbial functions across treatments (Fig. S6A). Various functional pathways associated with carbohydrate metabolism (starch degradation/biosynthesis, glycolysis, pentose phosphate pathway, glycogen degradation, gallate degradation), production of multiple exopolysaccharides (Colanic acid) membrane proteins (CDP-DAG), and defense response against various biotic and abiotic stress (gallate degradation, assimilatory sulfate reduction), etc. were found to be enriched in plants inoculated with *Xanthomonas*. In the absence of a dominant pathogen, the pathways related to biosynthesis and transport of amino acids that are functionally redundant for the growth (methionine biosynthesis, isoleucine biosynthesis), pathways associated with energy sources (Entner doudoroff, thiamine salvage), and defense pathways against plant defense (Isoleucine biosynthesis) were enriched (Fig. S6A). Upon O_3_ stress, the primary energy production source was the degradation of unsaturated fatty acids (beta-oxidation, pentose phosphate). Various defense-related pathways against oxygen stress and DNA repair (ubiquinol 7, pyrimidine (deoxy)nucleotides) and pathways related to oxygen-independent respiration (oxygen-independent heme b biosynthesis) were also found enriched (Fig. S6B). In the presence of both the pathogen and elevated ozone pathways related to purine nucleotide production and degradation was enriched (Fig. S6C).

### Microbial network topology is altered under combined pathogen and ozone stress

To assess whether pathogen infection and ozone stress alone or in combination affected overall microbial network complexity, we compared network topological features under abiotic, biotic, and combined stress against ambient and control conditions (Table S10A). While there were no statistically significant differences in overall global network properties, visual inspection of the bacterial network structures showed a distinct pattern of co-occurrence across various environmental conditions (Fig. 5). Microbial networks under ambient environment showed higher positive edge percentage, higher clustering coefficient, and lower average path length compared to elevated O_3_ (Table S10A). This suggests more positive interactions in ambient environments and that O_3_ stress may foster less complex and negative associations among community members. On the contrary, the presence of pathogen infection led to a more positive edge percentage, lower path length, higher modularity, and higher clustering coefficient, suggesting that all nodes were highly interlinked within the networks to form a more complex and stable network under pathogen stress (Table S10A). However, in the presence of combined biotic and abiotic stress, more positive interactions were found under ambient environment and control conditions with lower path length and higher clustering coefficient, suggesting that combined stress possibly creates less complex and less stable associations among community members (Table S10A).

**Fig. 5.**
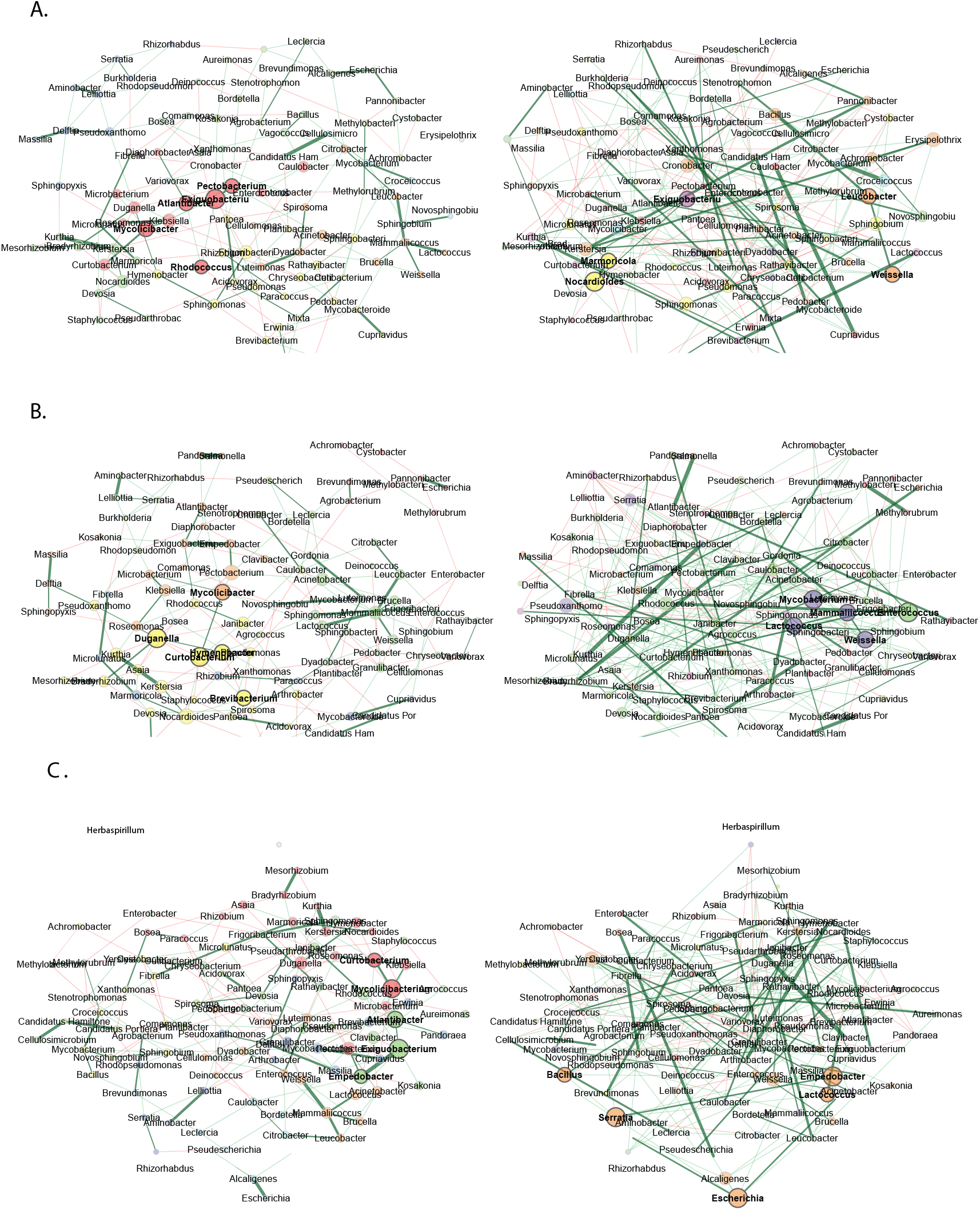
Comparison of bacterial association network across different environments. (A) Bacterial association network for the combined data set of ambient (left) and elevated O_3_ (right) in both cultivars under control conditions. (B) Bacterial association network for the combined data set from control (left) and inoculated (right) samples from both the cultivars under ambient environment. (C) Bacterial association network for the combined data set from control and ambient environment (left) and inoculated and elevated O_3_ (right) samples from both cultivars. Hub taxa are highlighted by bold text. Node color represents the cluster determined by greedy modularity optimization. Red edges correspond to negative correlations, while green edges correspond to positive correlations.

Next, the Jaccard index was used to test for similarities (Jacc□=□0, lowest similarity and Jacc□=□1, highest similarity) of the most central nodes in the networks under individual or combined stresses compared to the ambient environment. Degree, betweenness, and closeness centrality were found to be lower than expected by random for the comparison between all the treatment comparisons (Table S10B), suggesting significant differences in topology between these networks under individual or combined stresses compared to control. Regarding hub taxa, there was a significant difference among treatment groups when comparing control and pathogen stress or control and the ambient environment with combined stress (Table S10C). However, no shift in hub taxa was observed when comparing ozone stress to ambient conditions. Eigenvector centrality values for comparing networks under ambient vs. elevated ozone and control vs. inoculated samples suggested 25% dissimilarity in both networks (*p*□=□0.17), suggesting that despite the presence of abiotic or biotic stress, the most influential microbes in the network largely remain the same. In contrast, there was a significant change in the most influential microbes in the presence of combined stress (*p* < 0.001). Comparing the overall similarities of the two networks between the ambient vs individual stress or combined stress based on adjusted Rand index (ARI) indicated values greater than 0 (ARI=0.02, *p* = 0.07) for ambient vs O_3_ stress; control vs inoculated (ARI= 0.03, *p* = 0.02) and control and ambient environment vs combined stress (ARI = 0.10, *p* < 0.001) (Table S10D). These observations indicate that the partitions of species into communities show a low degree of similarity in these comparisons. These results, with differences in topology between these networks and dissimilarity in local network centrality measures, indicate that combination of biotic and abiotic stress results in a shift in the bacterial community interactions in the phyllosphere.

## Discussion

Plants respond to biotic and abiotic stresses via complex yet overlapping defense signaling pathways (57,58). These overlapping signaling pathways mean that climatic fluctuations can have a profound effect on the outcome of plant-pathogen interactions (59–61). As microbiome also plays a vital role in tolerance to biotic and abiotic stresses, the response of microbial communities to the challenge of individual or combined stresses through the processes of assembly, succession, interactions, and their associated functions may be a key factor in determining host susceptibility. In this study, we addressed this question using OTCs that allowed manipulating [O_3_] under natural field conditions and a temporal metagenomics approach.

While the apparent influence of elevated O_3_ was not observed on disease severity levels (measured as a percentage of spot symptoms and percentage of defoliation) on the susceptible cultivar, the resistant cultivar displayed higher disease severity under elevated O_3_ throughout the growing season as compared to ambient environment (Fig. 1). This increased disease severity on the resistant cultivar, however, was not translated into a significant difference in the relative abundance of pathogen *Xanthomonas* across these environmental conditions, partly attributing to the high variability in the relative abundance of *Xanthomonas* across three chambers. Indeed, such a culture-independent DNA sequencing method may not accurately indicate living pathogen cell count and may warrant confirmation of these findings with a culture-dependent pathogen population estimate. These results of higher disease severity in the resistant cultivar suggest a possible compromise in the resistance and pathogen population not likely being a determining factor for increased disease severity. Since the resistance in resistant pepper is governed by quantitative resistance loci that are less prone to compromise (62), various factors can lead to this increased susceptibility in resistant cultivars, such as altered pathogen aggressiveness due to changes in effector translocation. On the other hand, altered host immune response due to changes in plant hormone signaling as a preferred response of plants to abiotic stress depending on its timing and severity might also contribute towards enhanced disease severity on resistant cultivars. Specifically, previous work has shown that plants grown under abiotic stress conditions show increased disease susceptibility due to induction of the abscisic acid (ABA) pathway, which antagonizes the salicylic acid (SA) pathway involved in pathogen defense (63,64). Moreover, the oxidative damage of the plant cuticle caused by elevated [O_3_] might leave plants more exposed to pathogens, thus increasing disease severity (65). Higher disease severity in plants exposed to pathogens was also seen in O_3_ tolerant tomato (66) and spot blotch fungus in barley (67). Similar impacts of plant resistance to pathogen stress have been reported due to higher temperatures (60,68,69) and elevated CO_2_ (61). However, the outcome of plant-pathogen interactions in the presence of abiotic stress has been variable depending on the pathogen’s lifestyle, duration, timing, and severity of each biotic and abiotic stress (34,70).

Given the impact of altered [O3] resulting in a possible compromise in resistance to *Xanthomonas*, we evaluated the response of microbial communities in these altered environments. Like many phyllosphere studies, we observed an overall reduction in the prokaryotic and eukaryotic community richness and diversity throughout the growing season, as typical of seasonal succession. A significant decrease in richness and diversity observed with susceptible cultivar challenged with pathogens was not surprising as it has been observed in plants and humans involving diseases or inflammatory syndromes (Chen et al. 2020, Turpin et al. 2018). The phyllosphere microbiome assemblage was also dependent upon the host resistance (Fig. 2), which reflected in no significant reduction in richness and diversity of both prokaryotic and eukaryotic communities on resistant cultivar upon challenge with *Xanthomonas*. This result is in line with the findings of resistant maize having higher richness and diversity than the susceptible hybrids against southern and northern corn leaf blight (72,73). However, contrasting results were seen with higher bacterial richness and diversity in sugarcane cultivars susceptible to brown rust (74) and pumpkin against powdery mildew (75), suggesting interactions between host and pathogen can be an influential factor in shaping the plant microbiome in resistant cultivars. An increase in [O_3_] did not significantly affect microbial diversity and richness.

Taxonomic diversity across the two pepper cultivars during the growing season under various environmental conditions indicates Proteobacteria as the most abundant bacterial phylum, followed by Actinobacteria and Bacteroidetes (Fig S2). These bacterial families are ubiquitous members of the plant phyllosphere (1,76,77). Inoculation with the *Xanthomonas* resulted in fewer Firmicutes and Actinobacteria in both cultivars at the end season. Similar findings of disrupting the abundance of Firmicutes and Actinobacteria upon dysbiosis have been reported both in the phyllosphere and rhizosphere (71,78,79). Inoculation also significantly negatively affected the relative abundance of various genera, including *Pseudomonas, Pantoea*, and *Methylobacterium*. The negative interaction of these multiple taxa with *Xanthomonas* was in agreement with the previous findings in the tomato and pepper phyllosphere (38). Eukaryotic microbial communities constituted a low portion of the phyllosphere and were dominated mainly by genera *Meosziomyces, Golubevia, Paraphoma, Protomyces*, and *Cercospora*. These genera inhabit the phyllosphere and have been known for their pathogenic or biocontrol properties (80– 82).

Plant microbiome assembly starts as the seed is sown, and selection pressure exerted by various biotic and abiotic factors, cultivars, and dispersal events determines the microbiome diversity during the growing season (83,84). Phyllosphere microbes are simultaneously exposed to these various abiotic and abiotic stresses in the natural environment. Studies on plant’s response to a combination of abiotic and biotic stress have shown a unique and more complex response than that of individual stresses (85–87). The effect of combined stress is governed by various factors such as time, degree of stress, plant genotype, and other climatic or environmental factors, thus, not necessarily additive in nature (88). We asked how the phyllosphere microbiome would respond to individual and combined stress and contribute towards tolerance to these stresses. The presence of *Xanthomonas* was the primary driver of diversity in the pepper phyllosphere, followed by cultivar and time of sampling for the bacterial communities. In contrast, cultivar significantly influenced eukaryote diversity, followed by the time of sampling and the presence of *Xanthomonas*. There was a temporal influence of various factors in the microbial communities where cultivar significantly affected bacterial diversity during the mid-season, which then decreased during the end season, where the presence of *Xanthomonas* had a more substantial influence. Environment significantly shaped the eukaryote communities during the mid-season, while cultivars had more influence during the end season. Change in [O_3_] alone did not influence microbial communities associated with the susceptible cultivar. However, there was a shift in the community on resistant cultivars under elevated O_3_ in the absence of pathogen. The influence of host genotype and, thus, impact of host defense on microbial communities was evident in the presence of pathogen under control environment. Microbial communities on resistant cultivar under O_3_ stress alone were distinct compared to those observed in the presence of the pathogen alone, indicating that communities responded differently to each stress. Interestingly, the microbial communities associated with plants exposed to combined biotic and abiotic stress still preferred to respond similarly to that in the presence of a pathogen alone, regardless of the host genotype. It has been known that plants may prioritize one response over the other or use combination strategies to respond to multiple stress by activating antagonistic, additive, or completely unrelated molecular responses (89,90). Our result suggested that although the environment significantly affected microbial community composition in resistant cultivars, a dominant pathogen drives the community composition in the presence of combined stress. As microbial communities associated with the resistant cultivar were influenced by abiotic stress in the absence of a pathogen, we speculate that the change in microbial community structure could be because of the cultivar response towards the elevated O_3_ stress and that the resistance gene/s might also be involved in abiotic stress response pathway.

The functional diversity of microbial communities helps to understand the role of these microbial communities in a different environments (91). Overall microbial functional diversity remains insignificant, as measured by the abundance of microbial pathways across biotic, abiotic, and combined stress across two cultivars. This result of similar functional diversity across treatments suggests a functional redundancy across these diverse microorganisms in the phyllosphere in the presence of various biotic and abiotic stress. Functional redundancy in microbial communities is one of the ecological properties where diverse community members can perform the same functional role where a change in species diversity will have little impact on overall ecosystem functionality (92,93). The coexistence of taxonomically distant taxa capable of performing similar metabolic functions is widespread in the microbial system (94–97). Various metabolic pathways enriched under biotic, abiotic, and combined stress in both cultivars helps to understand the microbial environment experienced by the resident microbes. Our analysis of these pathways suggests that under elevated O_3,_ pathways related to beta-oxidation, oxygen-independent respiration, and pathways against oxygen stress and DNA repair were enriched. These pathways have been known to play an essential role in stress response in various microbial communities (98–101). Pathogen infection leads to the enrichment of various defense pathways. Upon pathogen infection pathways related to carbohydrate metabolism were enriched, a major pathway for energy and defense in bacteria (102–104). In the presence of combined stress, a single pathway related to purine nucleotide production and degradation was enriched. It has been shown that biosynthesis and degradation of purine nucleotides play an essential role in the growth and virulence of various fungal communities; microbial communities might be using these pathways for better growth and survival under combined stress (105,106). Microbial function in the ecosystem is determined not just because of the number and composition of taxa but because of the various positive, negative, direct, or indirect associations among the community members (107). In response to the pathogen challenge, we observed network parameters indicative of a densely connected network. These findings of enhanced positive and complex interaction among the microbial communities upon pathogen infection have been observed in both the phyllosphere and endosphere (108–110). Such densely connected network indicates cooperative interactions such as facilitation, mutualism or commensalism, and cross-feeding (111,112). Such connected networks, referred to as small-world networks (113), are hypothesized to harbor resistance toward disturbances. In contrast, microbial co-occurrence networks across abiotic stress and combined stress showed a similar trend of a relatively unstable random network compared to the control environment. This finding agrees with the notion that varying degrees of environmental stress disturb the stability of microbial communities in the soil (112). The observation from the similarity of the most central node suggests that microbial communities are considerably different across different treatments. The presence of a pathogen and combined stress considerably affected hub taxa. However, combined stress, but not individual stresses, had considerable influence on the most influential taxa, suggesting that plants respond to combined biotic and abiotic stress by changing the most influential microbial member in the random network. It would be interesting to dissect further the influence of individual genotypes and, thus influence of host defense responses on microbial community networks, as we observed a strong genotype effect on community composition. However, the present study is limited in sample size, which does not allow sufficient power to compare the network structure against two genotypes.

Overall, our study demonstrated that the microbial community responds to a change by not only altering community composition but also interactions among members and overall community function (Fig. 6). In our study, the microbial community response was primarily driven by pathogen as a major stress. Abiotic stress did have an impact on shifting the community composition with interactions among members indicative of a random network, although the most influential taxa remained the same. A combination of biotic and abiotic stresses resulted in a shift in the community like that observed in the presence of biotic stress. However, interactions among these community members were random and less connected, and the most influential taxa were distinct. This indicates that combined stress can cause destabilized microbial community structure. The extent to which such a destabilized community could impart functions such as stress tolerance or contribute towards an overall compromise in host susceptibility towards pathogens is unknown. However, this work provides a base for our understanding of the complex response of microbial communities and their interactions with the host genotype in response to a changing climate. As plants have evolved in association with their phyllosphere microbiome members, the community members identified in this study shown to be particularly susceptible to a shift in response to abiotic stress or combined stress might be crucial to evaluate for future work on harnessing the microbiome for stress-tolerant plants.

**Fig. 6.**
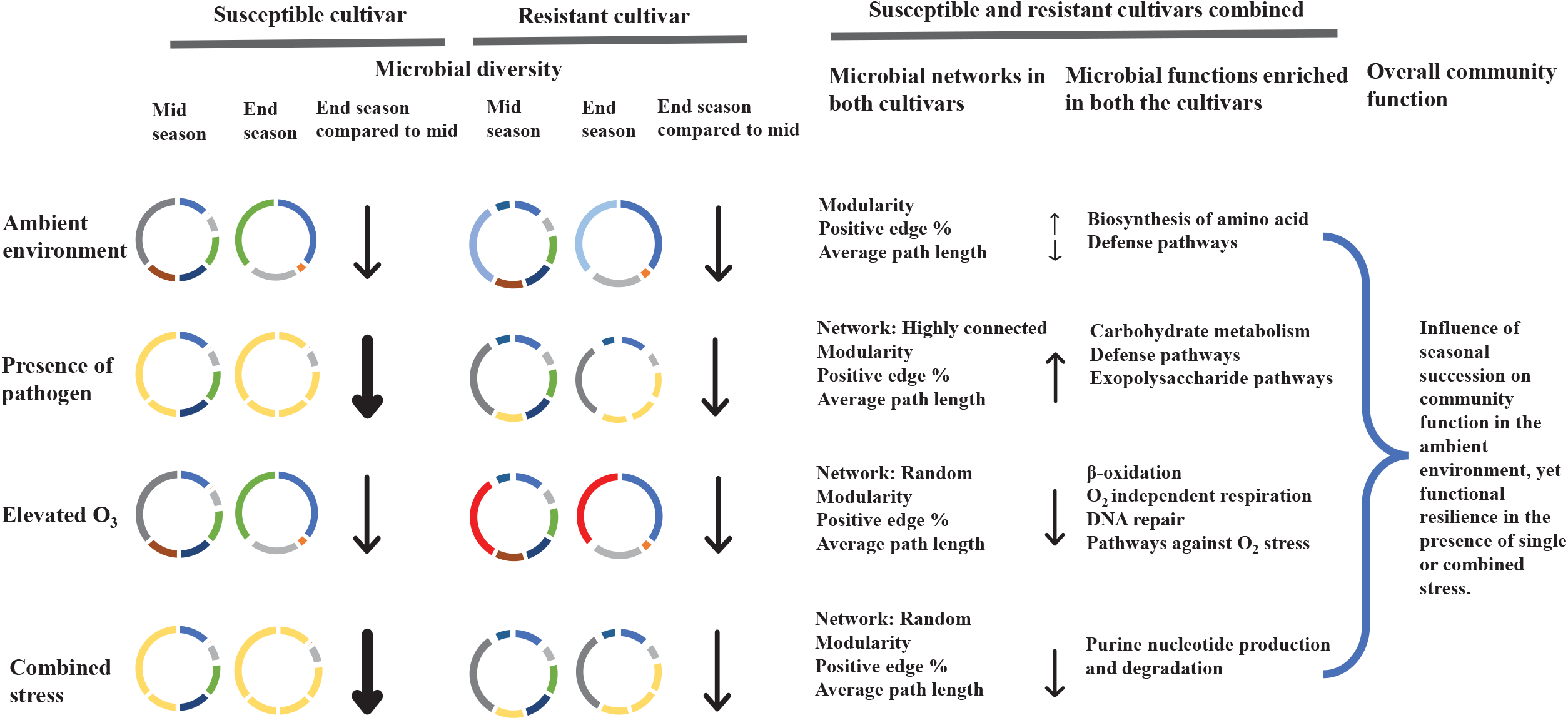
A schematic diagram highlighting the influence of different factors on phyllosphere microbial community structure and function. Different color donut charts represent the microbial community composition under different cultivars and treatments. The arrow represents the decrease or increase in response, and the weight of the arrow represents the magnitude of change.

## Data and code availability

Sequence data generated from this work have been deposited in the SRA (Sequencing Read Achieve) database under the BioProject accessions PRJNA889178. All other data and code used in this study are available in the following GitHub repository (https://github.com/Potnislab/AtDep_2021_metagenome).

## Acknowledgments

This work was supported by the Alabama Agricultural Experiment Station and the Hatch program (Project # 10108601) of the National Institute of Food and Agriculture, U.S. Department of Agriculture. We thank Auston Holland for his assistance in maintaining the plants in the greenhouse and throughout the experiment. We thank the members of Potnis, Leisner, and Sanz-Saez labs for their help with the initial planting. We thank Seth Johnston for setting up and maintaining the OTCs and fumigation. We thank Dr. David Young and the staff at the Alabama Supercomputer Authority for providing the computational resources necessary to conduct this work.

## Ethics declarations

### Competing interest

The author declares no competing interests.

---

Fig. S1. Study site and treatment design of Atmospheric Deposition Laboratory (AtDep) site at Auburn University. (A) Satellite image of AtDep site with individual chambers label where light blue circles are the chambers with the ambient environment and red circles are the chambers with elevated O_3_. Chambers 1-6 marked with yellow color are inoculated with *Xanthomonas perforans*, and chambers 7-12, with green color are control samples. (B) Individual open-top chamber at the AtDep site (4 × 5m). (C) Daily average ozone concentration in treatment chambers. Sampling points (mid and endseason) are marked by a red arrow.

Fig. S2. Stacked bar plots showing the relative abundance of dominant phylum across the samples for cultivar susceptible and resistant. Control samples are indicated by a green bar, while the yellow bar represents the inoculated samples. The time of sampling is indicated by Base (initial samples), Mid (mid season), and End (end season). Ambient represents the normal environment, whereas elevated O_3_ represents the chambers with elevated ozone in the treatment.

Fig. S3. Box plot showing the relative abundance of top 8 genera in cultivar susceptible across different treatments. Control samples are indicated by a green bar, while the yellow bar represents the inoculated samples. The blue box represents the samples taken during the midseason, while the orange box is the end-of-season samples. Different environmental conditions are represented by the ambient environment and elevated ozone under both inoculation and control treatments.

Fig. S4. Box plot showing the relative abundance of *Xanthomonas* across different treatments in resistant and susceptible cultivars during the mid and end season.

Fig. S5. Nonmetric Multidimensional Scaling (NMDS) ordination displaying diversity in community metabolic pathways across different treatment conditions in both the susceptible and resistant cultivars.

Fig. S6. Linear Discriminant Analysis Effect Size (LEfSe) of KEGG pathways between susceptible and resistant cultivar. Results were ranked by their Linear Discriminant Analysis (LDA) score. An FDR-adjusted *p*-value ≤ 0.05, as well as an LDA score ≥3, were used as thresholds to identify significant features. (A) pathways enriched in inoculated (yellow) and control (green), (B) pathways enriched in elevated ozone (red) and ambient environment (blue), (C) pathways enriched in combined stress (orange) and ambient environment (lime green).

## Notes

### Competing Interest Statement

The authors have declared no competing interest.

