## Supplementary material for "Biotic and abiotic stress distinctly drive the phyllosphere microbial community structure"

**This PDF file includes:**

Supplementary Results

Figure S1 to S6

**Other supplementary materials include: Tables S1-S10**

**Supplementary Results**

**Taxonomic profiling**

A temporal pattern was observed in that, *Actinobacteria* were the most abundant (~57%) in the base samples, and their abundance decreased over time both in inoculated (mid-season: 8 % and end season: 1%) and control plants (mid-season: 9 % and end season: 3%) shifting to Proteobacteria which was dominant both at the mid and end season samples (Figure S2). Phylum Bacteroidetes, although accounting for around 3% in the base samples, decreased in abundance to less than 1% at later time points and were negligible on the inoculated susceptible leaves. Phylum Firmicutes contributed to around 2% of the sequences during the mid-season. They were present in higher abundance under elevated $O_3$ conditions on the leaves of the control susceptible cultivar and in both inoculated and control plants in the resistant cultivar. This phylum was affected by inoculation on susceptible plants and was negligible across all the treatment conditions during the end season.

We next examined the deeper taxonomic ranks, specifically, genus, for further comparison of patterns affected by the presence of the pathogen and elevated $O_3$ or in combination. Susceptible control plants were dominated mainly by *Pseudomonas* (~5-60%), *Pantoea* (~3-28%), and *Sphingomonas* (3-15%), while *Xanthomonas* (~84-98%) and *Pseudomonas* (~1-4%) were the dominant genera in the inoculated plants. In resistant cultivar, *Pseudomonas* (6-40%), *Pantoea* (~2-44%), and *Methylobacterium* (~9-35%) were the dominant genera in control plants, while *Xanthomonas* (~2-87%), *Pseudomonas* (~2-27%), and *Methylobacterium* (~1-26%) were dominant on inoculated plants (Fig. 4A, Table S7). Among other genera contributing to the phyllosphere microbiome included *Pseudomonas*, *Pantoea*, *Methylobacterium*, *Sphingomonas*, *Methylorubrum*, *Stenotrophomonas,* and *Microbacterium*. The abundance of *Pseudomonas* increased over the growing season, except for resistant control plants, where abundance levels were higher during mid-season under elevated $O_3$ and stayed at similar levels throughout the growing season. *Pseudomonas* abundance was affected by inoculation, where a decrease in the abundance of *Pseudomonas* was proportional to the increase in *Xanthomonas* abundance on either cultivar. The mean relative abundance of *Pantoea* increased over time in control plants of either cultivar. The inoculation shifted the temporal pattern with a decrease in the mean relative abundance of *Pantoea* from mid to end season sampling. This decrease was statistically significant under elevated $O_3$ on inoculated samples of either cultivar (resistant cultivar: $p$ = 0.031; susceptible cultivar: $p$ = 0.007). Inoculation also had a slight effect on the mean relative abundance of *Methylobacterium* with a significant decrease with inoculation under ambient environment ($p$ = 0.005), under elevated $O_3$ in the resistant cultivar (Kruskal-Wallis, $p$ = 0.049), and ambient environment ($p$ = 0.022) in the susceptible cultivar. An increase in $O_3$ concentration has a negative effect on the *Methylobacterium* genus under control conditions and ambient environment (resistant cultivar: $p$ = 0.009, susceptible cultivar: $p$= 0.025). However, the environment did not affect the *Methylobacterium* genus in inoculated samples in both cultivars (Fig. S3).

Analysis of the eukaryotes diversity across our samples showed that the genera *Moesziomyces*, *Golubevia*, *Paraphoma, Protomyces,* and *Cercospora* were dominant across the samples (Fig. 4B, Table S8). Base samples were dominated by the genus *Pseudogymnoascus* (~64%), which was not observed during later time points. Genus *Bullera* had a higher relative abundance under elevated $O_3$ both in control (~ 32%) and inoculated plants (~16%) than that of ambient environment (~ 15% in control and 12% in inoculated) during the end season in both the cultivars. Inoculation resulted in lower *Zasmidium*, *Cladosporium*, *Aureobasidium*, *Alternaria,* and *Anthracocystis* genera in susceptible cultivars compared to resistant ones under both environmental conditions. Regardless of treatments, the relative abundance of the genus *Epicoccum* increased at the end season (~ 31%) compared to mid-season (~ 4%). In comparison, the genus *Protomyces* decreased during the end season (~ 3%) compared to mid-season (~ 11%). Inoculation did not affect the relative abundance of *Moesziomyces* in susceptible cultivars during both mid (~ 60%) and end season (~70%). In comparison, the presence of *Xanthomonas* had a negative effect on this genus during the end season (~ 12%) in resistant cultivars.

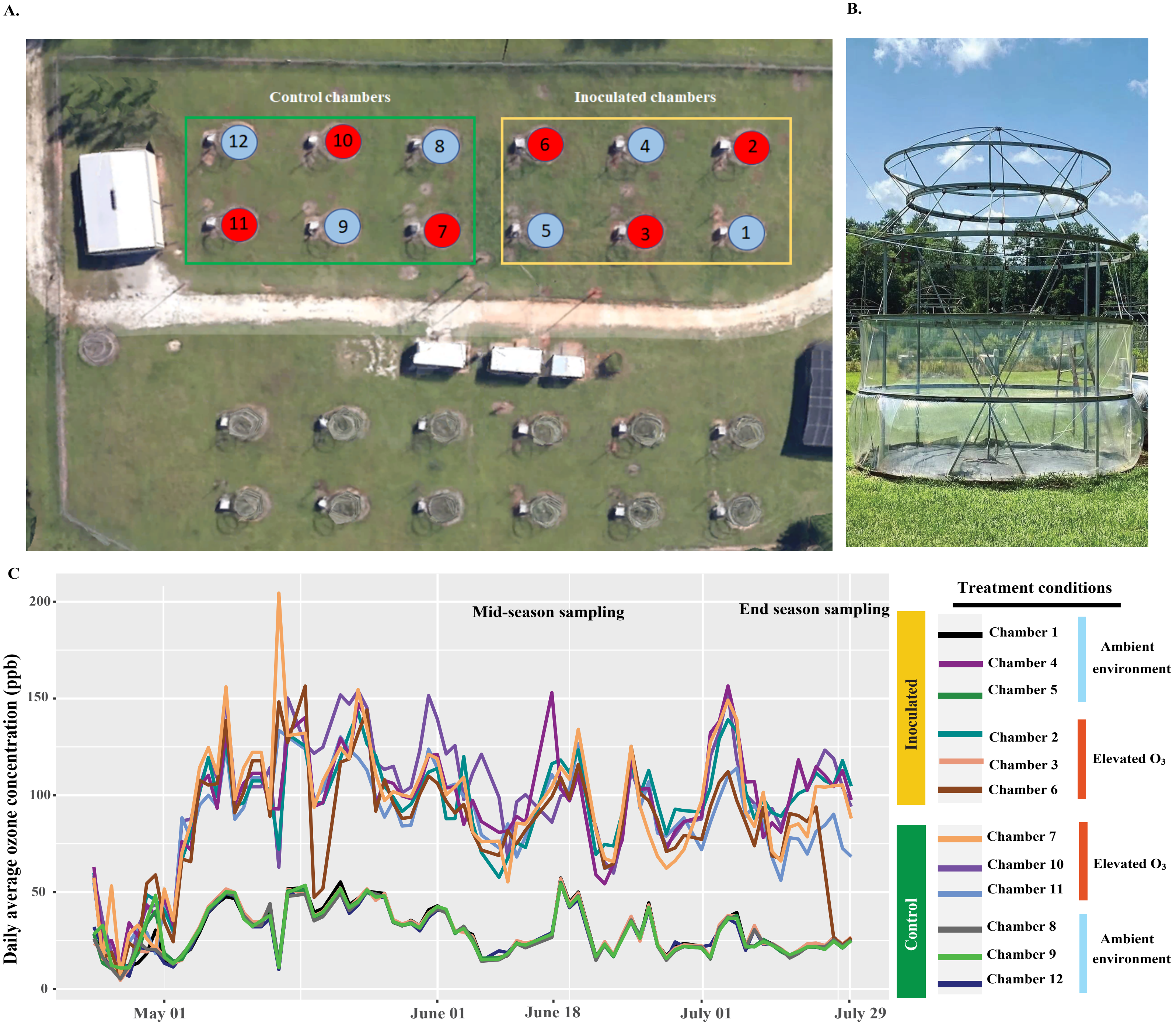


Fig. S1. Study site and treatment design of Atmospheric Deposition Laboratory (AtDep) site at Auburn University. (A) Satellite image of AtDep site with individual chambers label where light blue circles are the chambers with the ambient environment and red circles are the chambers with elevated $O_3$. Chambers 1-6 marked with yellow color are inoculated with *Xanthomonas perforans,* and chambers 7-12, with green color are control samples. (B) Individual open-top chamber at the AtDep site (4 x 5m). (C) Daily average ozone concentration in treatment chambers. Sampling points (mid and endseason) are marked by a red arrow.

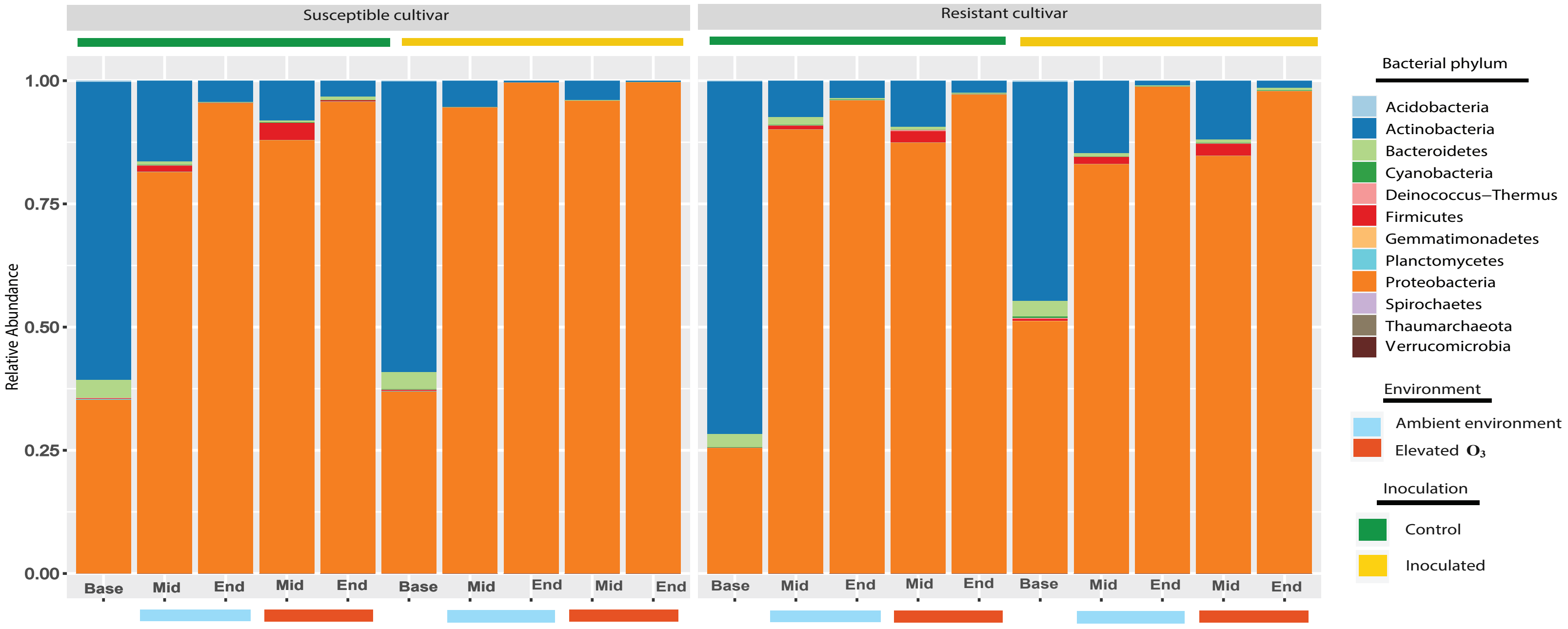


Fig. S2. Stacked bar plots showing the relative abundance of dominant phylum across the samples for cultivar susceptible and resistant. Control samples are indicated by a green bar, while the yellow bar represents the inoculated samples. The time of sampling is indicated by Base (initial samples), Mid (mid season), and End (end season). Ambient represents the normal environment, whereas elevated $O_3$ represents the chambers with elevated ozone in the treatment.

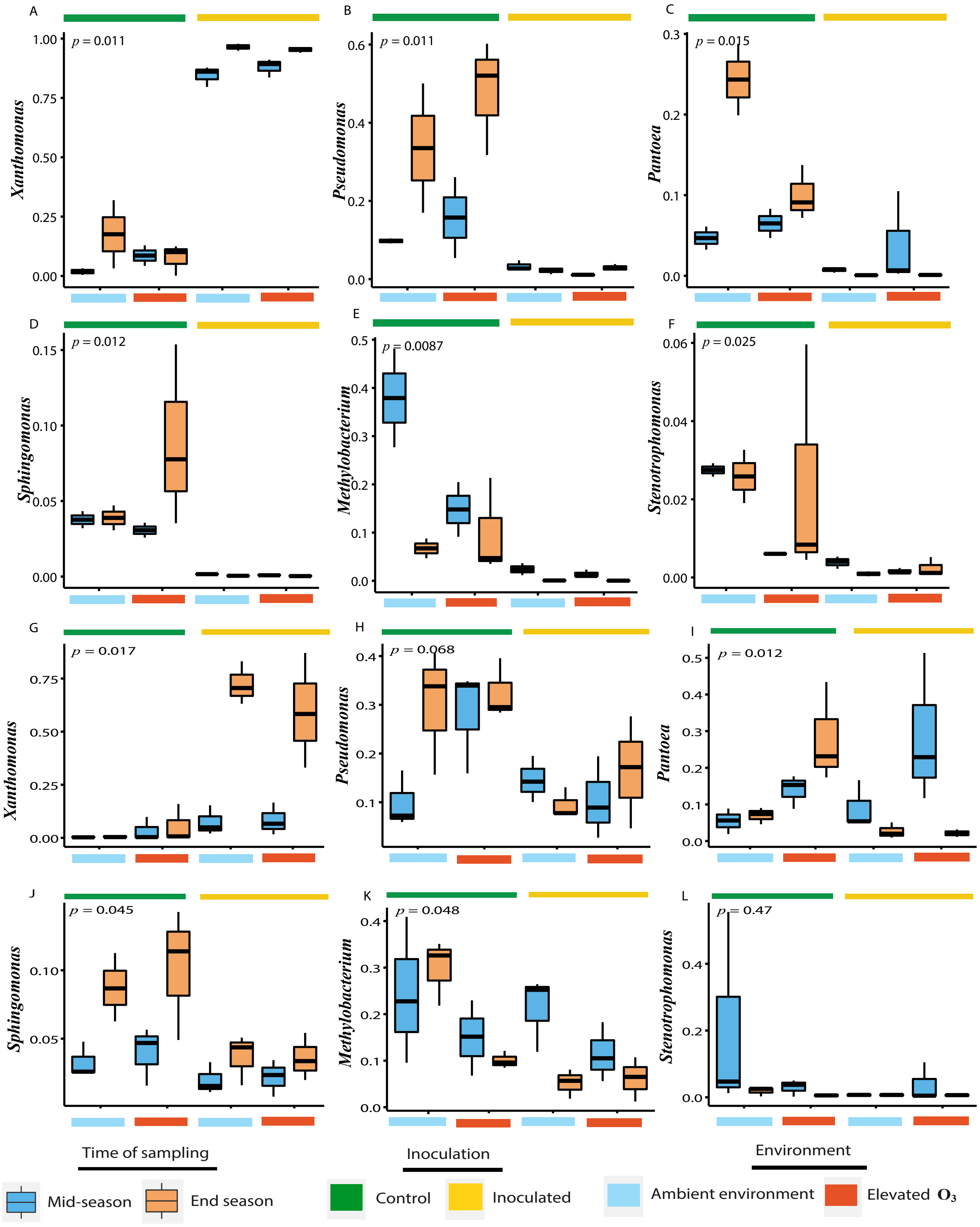


Fig. S3. Box plot showing the relative abundance of top 8 genera in cultivar susceptible across different treatments. Control samples are indicated by a green bar, while the yellow bar represents the inoculated samples. The blue box represents the samples taken during the midseason, while the orange box is the end-of-season samples. Different environmental conditions are represented by the ambient environment and elevated ozone under both inoculation and control treatments.

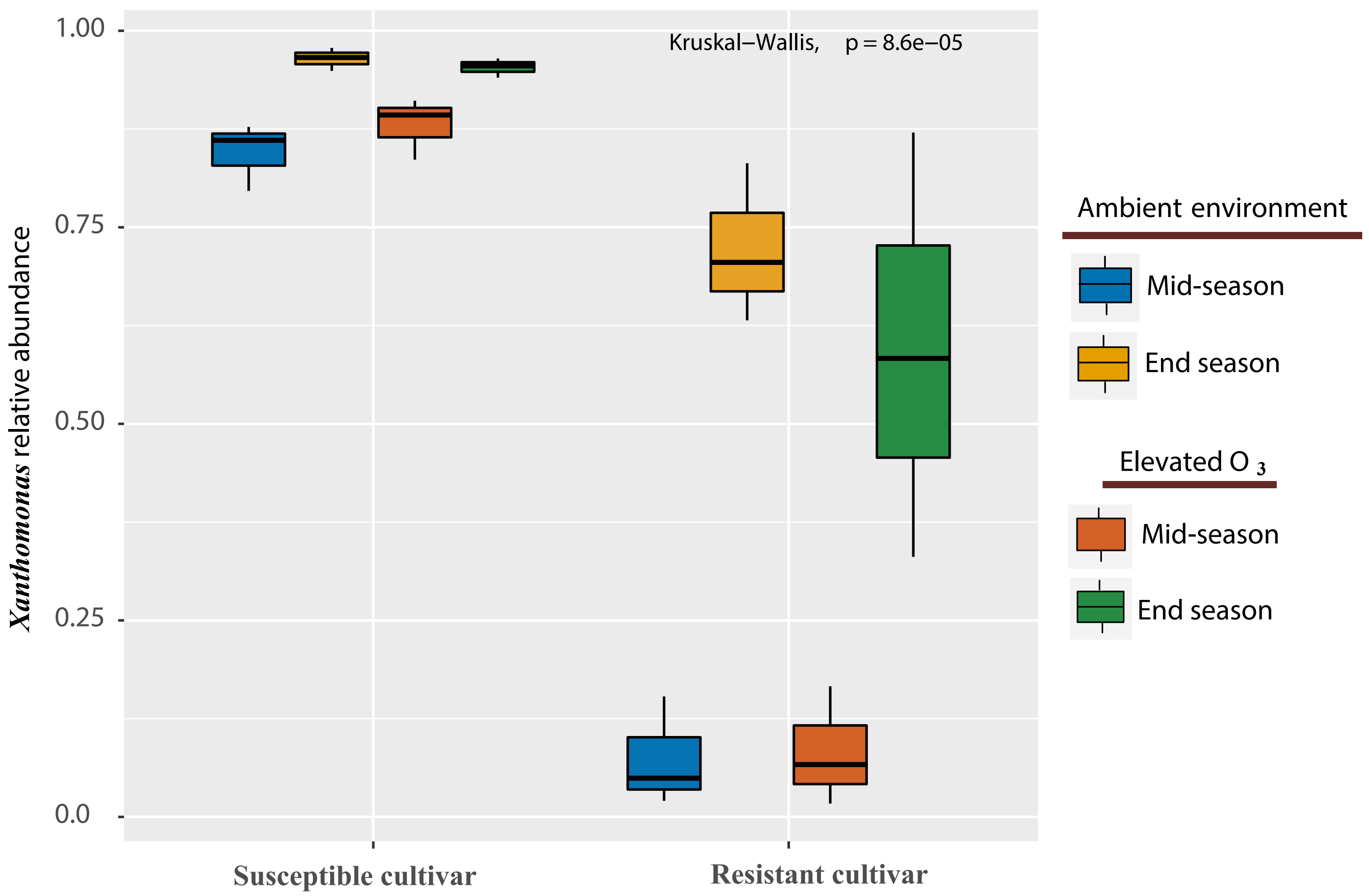


Fig. S4. Box plot showing the relative abundance of *Xanthomonas* across different treatments in resistant and susceptible cultivars during the mid and end season.

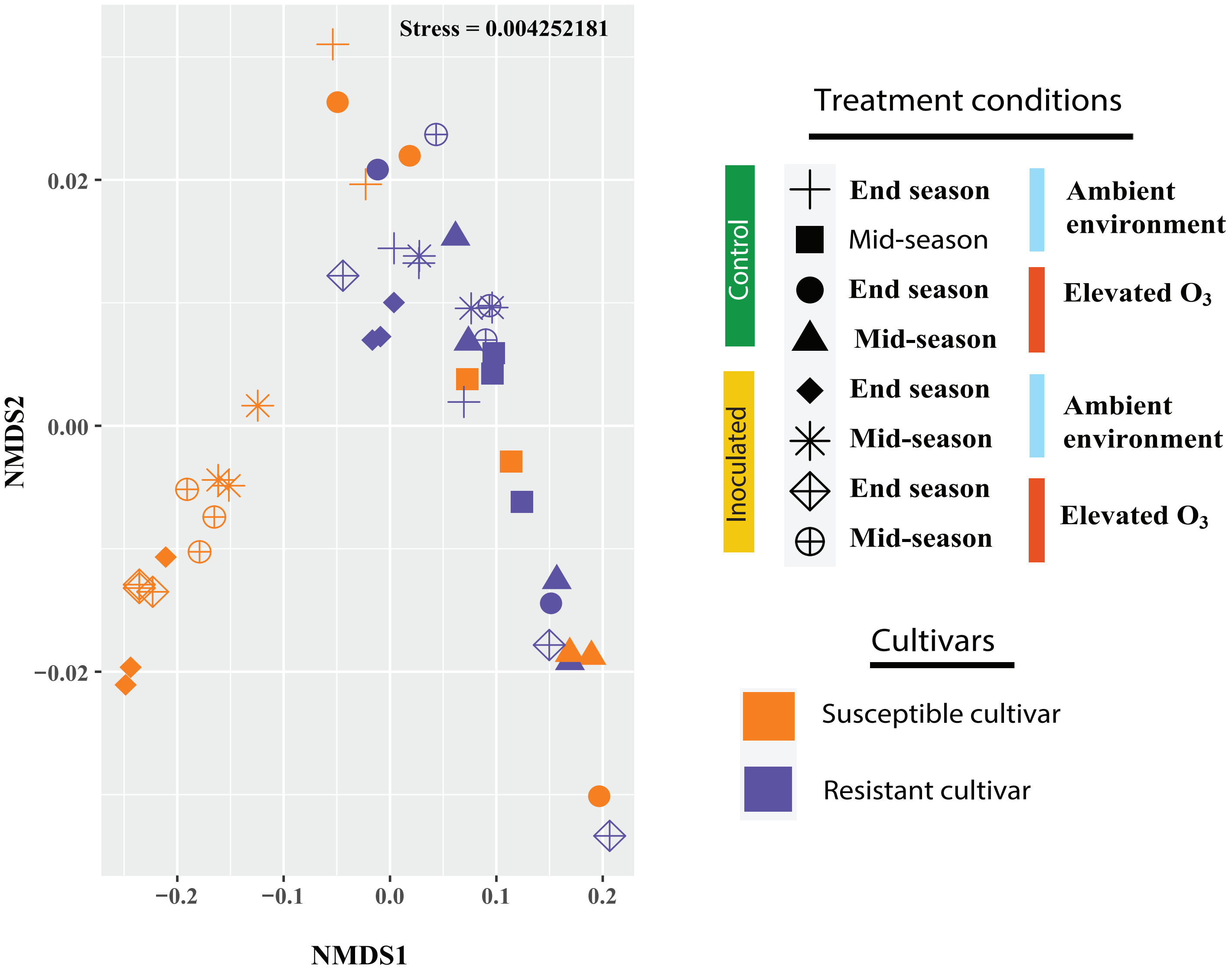


Fig. S5. Nonmetric Multidimensional Scaling (NMDS) ordination displaying diversity in community metabolic pathways across different treatment conditions in both the susceptible and resistant cultivars.

A.

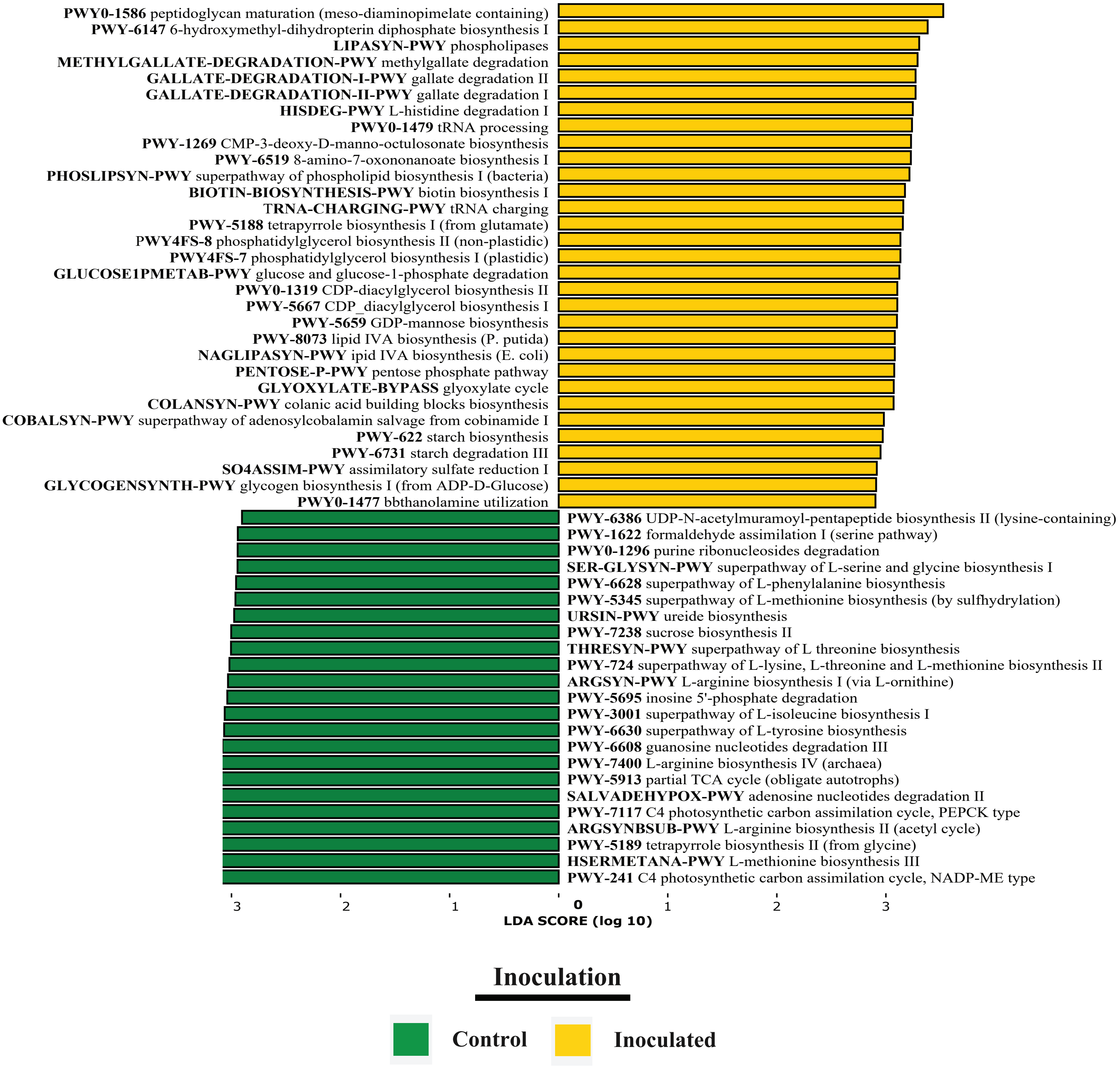
PWY0-1586 peptidoglycan maturation (meso-diaminopimelate containing)
PWY-6147 6-hydroxymethyl-dihydropterin diphosphate biosynthesis I
LIPASYN-PWY phospholipases
METHYLGALLATE-DEGRADATION-PWY methylgallate degradation
GALLATE-DEGRADATION-I-PWY gallate degradation II
GALLATE-DEGRADATION-II-PWY gallate degradation I
HISDEG-PWY L-histidine degradation I
PWY0-1479 tRNA processing
PWY-1269 CMP-3-deoxy-D-manno-octulosonate biosynthesis
PWY-6519 8-amino-7-oxononanoate biosynthesis I
PHOSLIPSYN-PWY superpathway of phospholipid biosynthesis I (bacteria)
BIOTIN-BIOSYNTHESIS-PWY biotin biosynthesis I
TRNA-CHARGING-PWY tRNA charging
PWY-5188 tetrapyrrole biosynthesis I (from glutamate)
PWY4FS-8 phosphatidylglycerol biosynthesis II (non-plastidic)
PWY4FS-7 phosphatidylglycerol biosynthesis I (plastidic)
GLUCOSE1PMETAB-PWY glucose and glucose-1-phosphate degradation
PWY0-1319 CDP-diacylglycerol biosynthesis II
PWY-5667 CDP_diacylglycerol biosynthesis I
PWY-5659 GDP-mannose biosynthesis
PWY-8073 lipid IVA biosynthesis (P. putida)
NAGLIPASYN-PWY ipid IVA biosynthesis (E. coli)
PENTOSE-P-PWY pentose phosphate pathway
GLYOXYLATE-BYPASS glyoxylate cycle
COLANSYN-PWY colanic acid building blocks biosynthesis
COBALSYN-PWY superpathway of adenosylcobalamin salvage from cobinamide I
PWY-622 starch biosynthesis
PWY-6731 starch degradation III
SO4ASSIM-PWY assimilatory sulfate reduction I
GLYCOGENSYNTH-PWY glycogen biosynthesis I (from ADP-D-Glucose)
PWY0-1477 bbthanolamine utilization
PWY-6386 UDP-N-acetylmuramoyl-pentapeptide biosynthesis II (lysine-containing)
PWY-1622 formaldehyde assimilation I (serine pathway)
PWY0-1296 purine ribonucleosides degradation
SER-GLYSYN-PWY superpathway of L-serine and glycine biosynthesis I
PWY-6628 superpathway of L-phenylalanine biosynthesis
PWY-5345 superpathway of L-methionine biosynthesis (by sulfhydrylation)
URSIN-PWY ureide biosynthesis
PWY-7238 sucrose biosynthesis II
THRESYN-PWY superpathway of L threonine biosynthesis
PWY-724 superpathway of L-lysine, L-threonine and L-methionine biosynthesis II
ARGSYN-PWY L-arginine biosynthesis I (via L-ornithine)
PWY-5695 inosine 5'-phosphate degradation
PWY-3001 superpathway of L-isoleucine biosynthesis I
PWY-6630 superpathway of L-tyrosine biosynthesis
PWY-6608 guanosine nucleotides degradation III
PWY-7400 L-arginine biosynthesis IV (archaea)
PWY-5913 partial TCA cycle (obligate autotrophs)
SALVADEHYPOX-PWY adenosine nucleotides degradation II
PWY-7117 C4 photosynthetic carbon assimilation cycle, PEPCK type
ARGSYNBSUB-PWY L-arginine biosynthesis II (acetyl cycle)
PWY-5189 tetrapyrrole biosynthesis II (from glycine)
HSERMETANA-PWY L-methionine biosynthesis III
PWY-241 C4 photosynthetic carbon assimilation cycle, NADP-ME type
3
2
1
0
1
2
3
LDA SCORE (log 10)
Inoculation
Control
Inoculated

B.

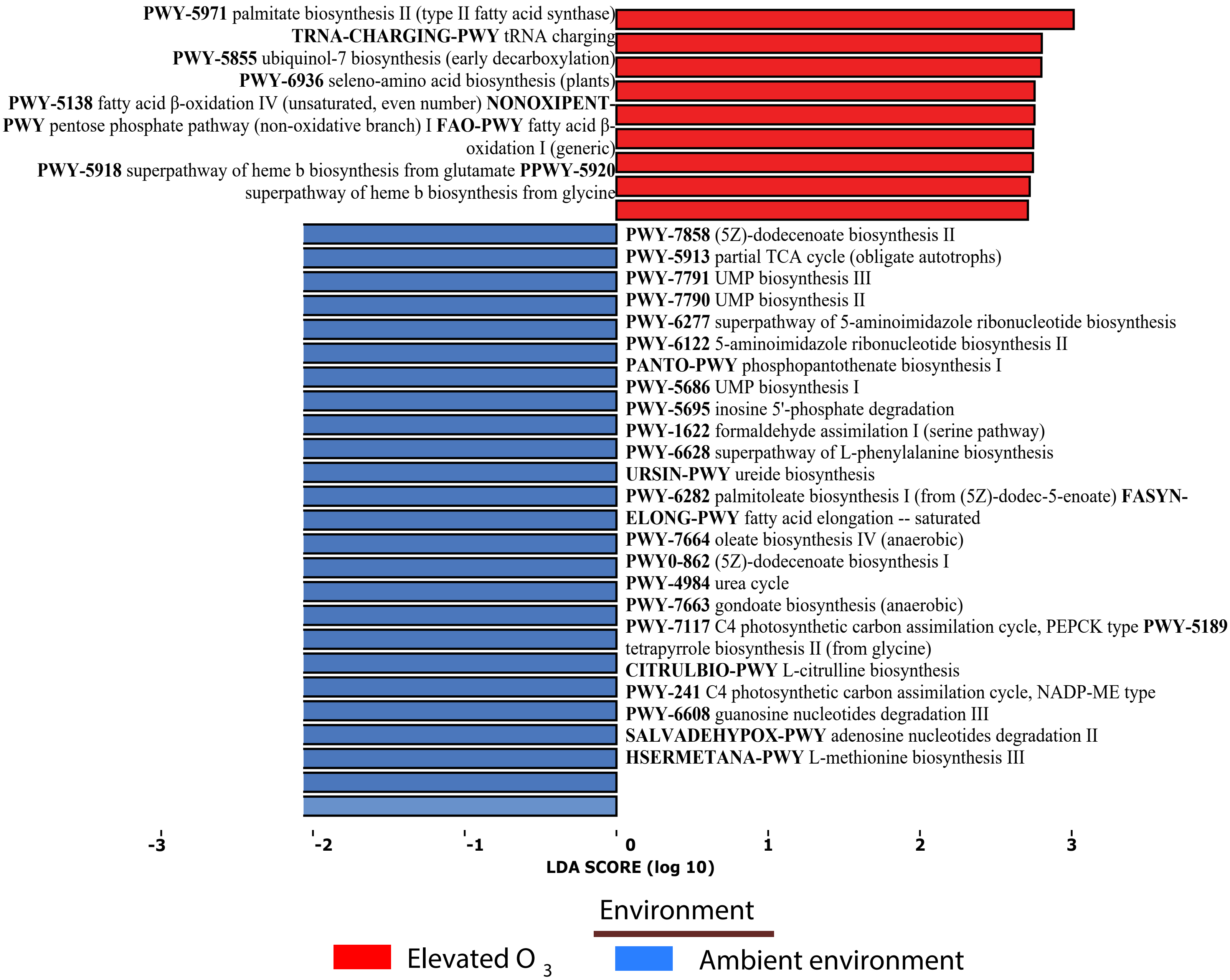


C.

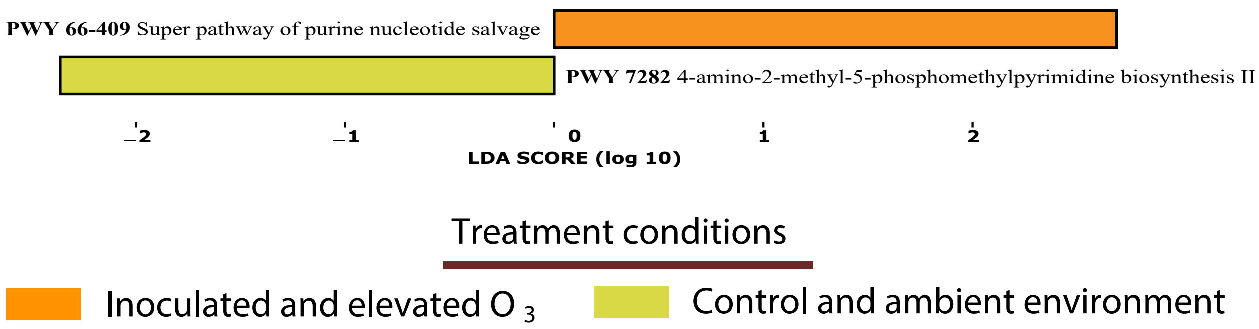


Fig. S6. Linear Discriminant Analysis Effect Size (LEfSe) of KEGG pathways between susceptible and resistant cultivar. Results were ranked by their Linear Discriminant Analysis (LDA) score. An FDR-adjusted $p$-value ≤ 0.05, as well as an LDA score ≥3, were used as thresholds to identify significant features. (A) pathways enriched in inoculated (yellow) and control (green), (B) pathways enriched in elevated ozone (red) and ambient environment (blue), (C) pathways enriched in combined stress (orange) and ambient environment (lime green).
